# Changing pollinator communities along a disturbance gradient in the Sundarbans mangrove forest: a case study on *Acanthus ilicifolius* and *Avicennia officinalis*

**DOI:** 10.1101/2020.02.20.953166

**Authors:** Asma Akter, Paolo Biella, Péter Batáry, Jan Klečka

## Abstract

The Sundarbans, the largest mangrove forest in the world and a UNESCO world heritage site has been facing an increasing pressure of habitat destruction. Yet, no study has been conducted to test how human disturbances are affecting plant-pollinator interactions in this unique ecosystem. Hence, we aimed to provide the first insight of the impact of habitat loss and human disturbances on the pollinator communities in the Sundarbans. We selected 12 sites in the North-Western region of the Sundarbans, along a gradient of decreasing habitat loss and human activities from forest fragments near human settlements to continuous pristine forest, where we studied insect pollinators of two mangrove plant species, *Acanthus ilicifolius* and *Avicennia officinalis*. Our results show that different pollinator groups responded to the disturbance gradient differently. For example, the abundance of *Apis dorsata, one of the three local species of honey bees*, increased gradually from the village area towards the deep pristine forest. On the other hand, *A. cerana* and *A. florea* were found in the village sites and completely absent in the deep forest. Although pollinator community composition changed along the disturbance gradient, their efficacy in pollination did not seem to be significantly affected. However, lower plant diversity and low understory plant cover in the forest patches nearby the village indicated that human disturbances not only affected pollinator community composition but also played a major negative role in the regeneration of the forest. Our study provides first insights into plant-pollinator interactions in the Sundarbans and demonstrates that more research is needed to inform conservation of this unique habitat.

## 1. Introduction

Human destruction of natural habitats and alteration of landscapes are considered as major drivers of the world-wide forest loss and fragmentation (Aizen and Feinsinger, 1994; Fischer and David, 2007). This increasing disturbance and habitat loss not only change the distribution and abundance of different organisms but also affect species interactions, which may be amplified into long-term effects on the forest ecosystem (Fortuna and Bascompte, 2006). Plant-pollinator interactions play a crucial role in ecosystem function as around 90% of angiosperm species rely on pollinators at least to some extent for their sexual reproduction (Ollerton *et al*. 2011, Potts *et al*. 2016). This makes pollinators an essential component to maintain biodiversity and ecosystem integrity (Kearns *et al*. 1998; Potts *et al*. 2003).

In most forest ecosystems, the fringe of forest is generally under pressure of high human activities, e.g. illegal collection of wood for fuel, house building materials, and agricultural tools along with regular grazing of domestic animals. These frequent disturbances affect the forest structure and interrupt the ability of the understory species to regenerate (Smiet, 1992). Alterations of natural habitats can affect plant-pollinator interactions in different ways. On the one hand, pollinators can be affected by the lack of suitable habitat and resources, which may determine their performance (Ward and Johnson, 2005). From the pollinators’ perspective, destruction of habitats or reduction in the availability of food (nectar and pollen) and nesting sites are expected to reduce species richness, abundance and homogenize species composition (Sameiima *et al*., 2004, Steffan-Dewenter & Westphal, 2008, Biella *et al*. 2019, in press). Furthermore, increased flight distance among habitat fragments can cause less effective pollen transfer (Aizen and Harder 2007). Pollinator abundance can also decrease due to lower attractiveness of isolated fragments, small population size, or reduced density of flowering plants (Cheptou and Avendano, 2006). Consequently, plants may suffer reduced seed set (Ward and Johnson, 2005). Overall, the stability of plant-pollinator interactions tends to be altered when native habitat is changed or removed. Even small disturbances may cause disruption of plant–pollinator interactions within the remaining habitat patches in fragmented landscapes (Keitt, 2009). Plant’s evolutionary dependence on pollinator communities for the pollination and reproduction increases the susceptibility to habitat loss and human disturbances and in return, pollinator diversity, abundance and foraging behaviour might also get affected as a consequence (Quesada *et al*., 2011). However, different pollinator communities may react to the forest loss and human disturbances at different scales and depend on the flower composition and environmental conditions both at the local and landscape scales (Hamer *et al*., 2000; Breitbach *et al*., 2012).

Unlike most terrestrial ecosystems, mangroves are naturally fragmented, architecturally simple and often have limited species diversity, but with a number of uniquely adapted species (Vannucci, 2001, Alongi, 2002). Heavily populated coastal zones have accelerated the widespread clearing of mangroves for coastal development, aquaculture, or other resource uses (Polidoro *et al*., 2010) and have led to further forest destruction, fragmentation and habitat loss. Globally, around 20%- 35% of mangroves have been lost since the 1980s and approximately 1% of the mangrove areas are disappearing per year (Valiela *et al*. 2001; FAO, 2003; FAO, 2007). An extreme example of forest loss and habitat destruction is the Sundarbans mangrove forest, situated in the south-western Bangladesh, which is the world’s largest continuous mangrove forest (Sarkar *et al*., 2016). Nearly 50% of the forest has been lost since the 1950s because of inadequate habitat protection, and large-scale habitat alteration (Feller *et al*., 2010). Historical human pressures have severely degraded the Sundarbans ecosystem by depleting forest tree stock (Ellison *et al*., 2000) and causing habitat loss. While natural disturbances determine both regional and global forest dynamics and diversity (Masaki *et al*., 1999; Sheil, 1999), anthropogenic activities may locally regulate the regeneration dynamics of forests and influencing the structure and floristic composition of the lowland forest (Horn and Hickey, 1991). A recent study by Sarkar *et al*., (2019) also stated an increasing trend of compositional homogeneity in the plant diversity and radical shifts in species composition in the Sundarbans. Introduction of non-mangrove plants in the forest, either intentionally or accidentally, increasing population of invasive plant species, decreasing population of certain mangrove plant species (Sarkar *et al*., 2019) and keeping honeybees (mainly *Apis cerana*) for apiculture along the forest edge for honey production are also sources of concern and their impact on this forest must be assessed to maintain local biodiversity. Despite the numerous ecosystem services provided by this mangrove forests (Walters *et al*., 2008), very little is known about pollinator communities of this forest (Pandit and Choudhury, 2001; Hermansen, *et al*., 2014), and there have been no studies evaluating the impact of human disturbances and habitat loss on the pollinator communities, their interactions with local plants and plant reproduction.

While studies on the pollination ecology and biology of mangrove plants around the world are frequent (Aluri, 2019), studies on the pollinator communities and pollination efficacy in the Sundarbans are scarce. Only a few studies focused on the pollinator communities of the Indian part of the Sundarbans (Mitra *et al*., 2015; Chakraborti *et al*., 2019). Generally, *Apis dorsata* is considered to be the most common pollinating insects in the Sundarbans (Gani, 2001; Mitra *et al*., 2015; Chakraborti *et al*., 2019), especially in the major flowering season (from March to June), while other *Apis* species and solitary bees are also common in this forest and in other mangrove forests in the Indian subcontinent. Here, we targeted two plants species, *Acanthus ilicifolius* and *Avicennia officinalis*, to compare the pollinator communities along the disturbance gradient in the Sundarbans and test the impact of the disturbances on the plant-pollinator communities and pollination. Reproduction bilogy of these two species is well known (Aluri *et al*., 1994; Aluri *et al*., 2012; Aluri *et al*., 2017), although possible effects of anthropogenic activities and disturbances on their reproduction have not yet been studied. However, such studies are essential to predict the sustainability of a forest ecosystem and primary requirement to take any conservation decision. Therefore, we addressed four questions: i) Does the plant diversity and abundance of floral resources decrease with the increasing human disturbances? ii) Does the abundance of flower visitors decrease and the composition of their community change along the gradient of human impact? iii) Do differences in pollinator visitation along the gradient affect the level of pollination and seed production of selected plants, with seed set reduced in disturbed sites? iv) And, what kind of conservation measures should be taken to protect both plant and pollinator communities?

## 2. Methods

### 2.1 Study area

This study took place in the North Eastern part of Sundarban Mangrove Forest in Bangladesh, located nearby Munshigang, Shyamnagar, Satkhira (N 22°16’78, E 89°11’58). The Sundarbans is protected as a UNESCO world heritage site. There are three protected sites in this forest in the Bangladesh sites of the Sundarbans: East Wildlife Sanctuary (ES, 312 km2), South Wildlife Sanctuary (SS, 370 km2), and West Wildlife Sanctuary (WS, 715 km2) (Gopal and Chauhan, 2006).

However, the part of the forest we studied is outside of these protected areas and highly disturbed by human activities and facing a high rate of biodiversity loss. The forest is distinctly isolated by the river ‘Pankhali’ from the adjacent human settlements, though fragmented forest patches are still found inside the village areas.

Based on the distance of isolated forest patches from the forest, canopy and ground cover and intensity of human disturbances, we selected twelve sites (Fig. 1C, site characteristics: supplementary table 1). Therefore, our study sites expanded from the most fragmented and isolated forest patches in the village to the pristine forest sections and from the most to the least affected by anthropogenic activities. The maximum distance from the most disturbed site to the least disturbed site was ca. 10 km. Forest patches inside the village were adjacent to the high-density human settlement and completely exposed to their daily life activities. The grounds of these sites had no or very little understory vegetation and distance from the continuous forest was 1-2.5 km. Forest patches which were close to the river, were also exposed to high human activities and had little understory vegetation as well. On the other hand, sites on the opposite side of the river in the continuous forest with moderate human impact had around 50 % ground covered by understory plants. Finally, forest patches which were deep in the forest and the farthest from the village, were least or not disturbed at all, high in plant density and almost fully covered by the herbaceous and shrub plants (Data: https://doi.org/10.6084/m9.figshare.11877615). This part of the forest is only occasionally visited by the forest department for regular security checking and by the honey-collectors from wild *Apis dorsata* colonies, thus has the lowest human disturbance.

**Table 1:**
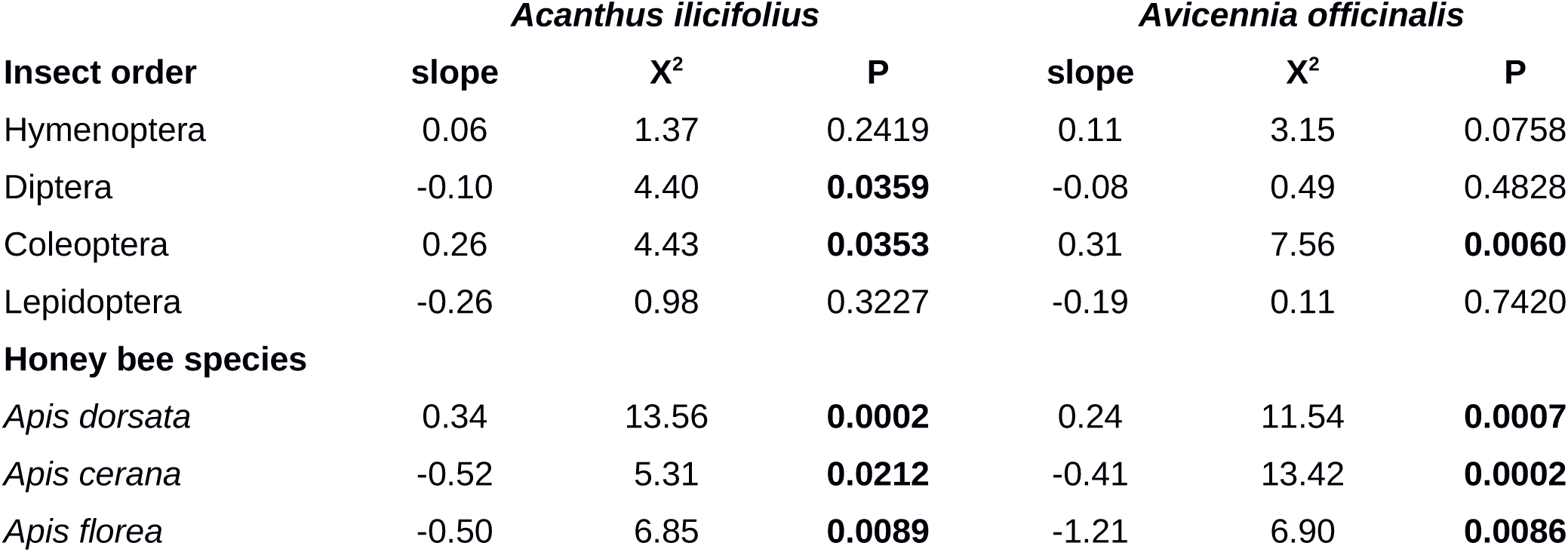
Visitation rate by different groups of pollinators on the two plant species. Results of statistical tests (GLMM) of the changes of the visitation rate by insect orders and individual species of honey bees along the disturbance gradient from the village towards the forest interior. Likelihood ratio test was used to test the statistical significance of each fitted relationship.

**Fig 1:**
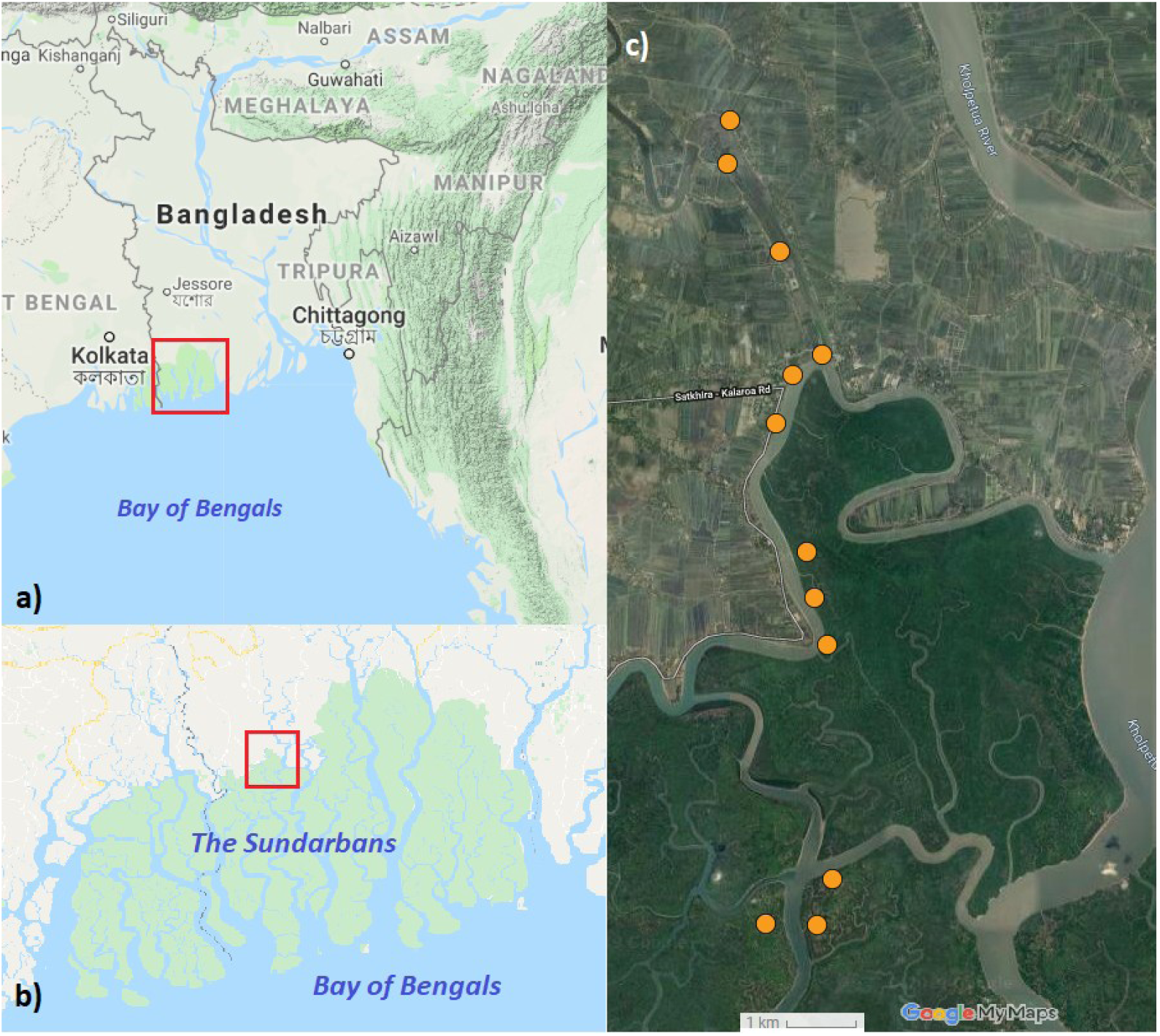
Location of the study area and the position of individual sites. Location of the Sundarbans mangrove forest in Bangladesh (A.) and location of our study area at the inland edge of the mangrove forest (B.). Location of individual sampling sites in the village and forest area (C.). Map data: Google, Imagery: TerraMetrics.

**Figure 2:**
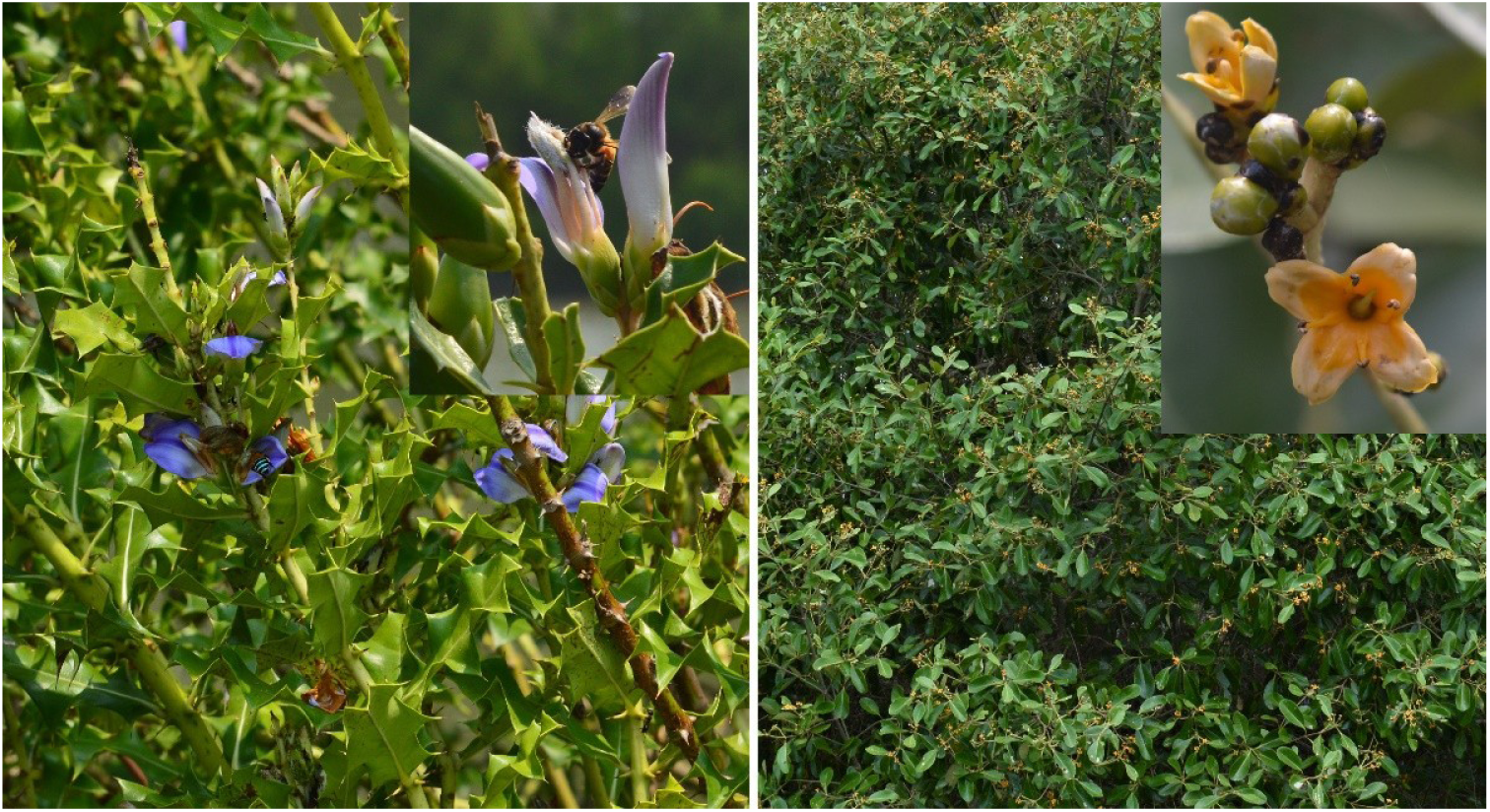
The target plant species of our study. *Acanthus ilicifolius* plant and flower, being visited by *Apis dorsata* (left); *Avicennia officinalis* plant and flower (right).

### 2.2 Plant diversity assessment

We surveyed the vegetation of all sites during our fieldwork. We identified all the species in each site and estimated total canopy cover, cover of the understory layer, and percentage cover by individual plant species. Only two plant species were flowering at all sites during the time of our study. Hence, these two species were chosen for a detailed study.

#### 2.2.1. *Acanthus ilicifollius* L

(Lamiales: Acanthaceae) is an evergreen, non-viviparous, semi-woody spiny shrub, which grows up to two meters. It has a wide range of distribution; it occurs from Western India through the North-Eastern China to Southern Australia (Tomlinson, 1986). It is commonly found along the edges of estuaries, canals and river banks. In our study site, they were found also found in the interior of the forest as only those forest patches were chosen which were flooded by the tidal flow and go underwater (Photos in the supplementary documents, S1). This species is very important for the accumulation of soil sediments and stabilization of the ground in brackish water areas. Inflorescence is spike, terminal, flower is large, showy, light blue to purple coloured, contains one large petal, four stamens, is bisexual and semi-tubular in shape (Aluri *et al*, 2017). The species produces nectar and pollen, has a mixed breeding system where out-crossing plays the most important role and it was reported to be pollinated by large bees (Aluri, 1990). Flowering time in the Sundarbans spans from April to June but can be different for other parts of its distribution zone (Ramasubramanian *et al*., 2003, Upadhyay and Mishra, 2010). Fruit is a capsule containing up to four seeds that disperse by effective anemochory especially during the dry season (Aluri *et al*., 2017).

#### 2.2.2 *Avicennia officinalis* L

(Lamiales: Acanthaceae) is a common viviparous mangrove tree, which has a wide range of distribution from Southern India through Indo-Malaya, to New Guinea and the Eastern Australia (Tomlinson, 1986, Duke 1991). *A. officinalis* can tolerate a wide range of salinity and occurs dominantly in soils with high salinity, and frequent and long duration of tidal inundation, although their abundance is higher towards the landward sites in the Sundarbans (Joshi and Ghose, 2003). It is a medium-sized tree, typically twenty meters tall, but can be up to thirty meters tall and contains pneumatophores. Inflorescence is spike, flower is small, yet the largest among the *Avicennia* species, orange-yellow coloured with four petals, four stamens, bisexual, open (Aluri, *et al*., 2012). It produces both nectar and pollen, is self-compatible although it is protandric, has a long flowering period suggesting its adaptation for cross-pollination, and is mostly pollinated by bees and flies (Aluri, *et al*., 2012). Flowering time is from April to August, depending on the location. Flowering is triggered by the rain and may vary even over a short distance (Opler *et al*, 1976; Reddy *et al*., 1995). Like other *Avicennia spp*., *A. officinalis* contains 4 ovules but in general only one ovule develops into mature seed, which is non-dormant and germinates while the fruit is still attached to the tree, thus are crypto-viviparous in character (Tomlinson, 1986, Aluri, *et al*., 2012).

### 2.3 Insect sampling

We observed and sampled flower-visiting insects form the two locally most abundant plant species, *Acanthus ilicifolius* and *Avicennia officinalis*. We surveyed them in May-June 2018, during the peak flowering time. We conducted our observations and sampled floral visitors for ten days. In each site, we sampled for 20-30 minutes in each session, replicated six times, which resulted into 120-160 minutes of observation for each species per site. During the high tide, a vast area of the forest is flooded, which restricted our fieldwork to 4-6 hours per day (Data: https://doi.org/10.6084/m9.figshare.11877615). For our observations we set up three collection windows with a size of 1 m^2^ for each plant species and we always sampled in the same windows. For both species, we counted the number of inflorescences and number of flowers per inflorescence within the window. For the *Avicennia officinalis*, we also measured the total flower cover in each window. We had three windows in every site for both species. Insects were observed and collected by netting, from 7 a.m. to 5 p.m., in sunny and warm condition, no observation was made under rain, storm or high winds. We determined the flower visitors when they touched the reproductive parts of the flower or entered the flower with a tubular shape. The three honey bee species and some other conspicuous flower visitors were released after counting, as they were easily recognizable. The rest of the captured insect were stored in the freezer after the collection and later mounted and stored dry in boxes for identification. Insects were identified by the authors and experts by using their expertise and various identification keys and taxonomical revisions of individual genera (Brunetti, 1923, Curran, 1947; Kumar & Sharma, 2015; Goulet & Huber, 1993; Pesenko & Pauly, 2005, Schmid-Egger, 2011). Bees, wasps and hoverflies were identified at the species or genus level, while other insects were identified up to family and only in few cases up to superfamily. We used the concept of ‘morphospecies’ denoted as sp.1, sp.2 etc. when species-level identification was not possible.

### 2.4 Pistil collection and pollen tubes analysis

In order to measure the pollination efficacy by the pollinators in different sites, we counted the number of pollen tubes in pistils as a proxy to pollen deposition. Although counting the number of pollen tubes does not differentiate between self- and cross-pollination, the number of pollen tubes growing in pistils is linked to the deposition of viable conspecific pollen and to seed production, hence provides information about pollination efficacy (Alonso *et al*., 2012, Biellla *et al*., 2019). We collected pistils at the end of our fieldwork to determine the impact of pollinator efficacy for each plant. Pistils were collected from 30 flowers per site excluding the plants where pollinators were observed and only form those flowers where female phase was over and stigmas were no longer receptive for pollen. Collected stigmas were stored in Formalin-Acetic-Acid solution (FAA) at room temperature. To assess the pollen tube growth, pistils were softened and stained by following the technique of Martin (1959). Pistils of both species were softened in 1M NAOH for 24 hours. After softening, they were stained with 0.1% Aniline blue in 0.1M K2HPO4 for 15 hours in the dark. After completing staining, pistils were washed and mounted in 50% Glycerine drop on glass slides, flattened evenly and covered with coverslips for observing under the fluorescence microscope and counted (Fig. 3). All the processes were done at room temperature and after the observations, samples were stored at 4°C for future reference.

**Figure 3:**
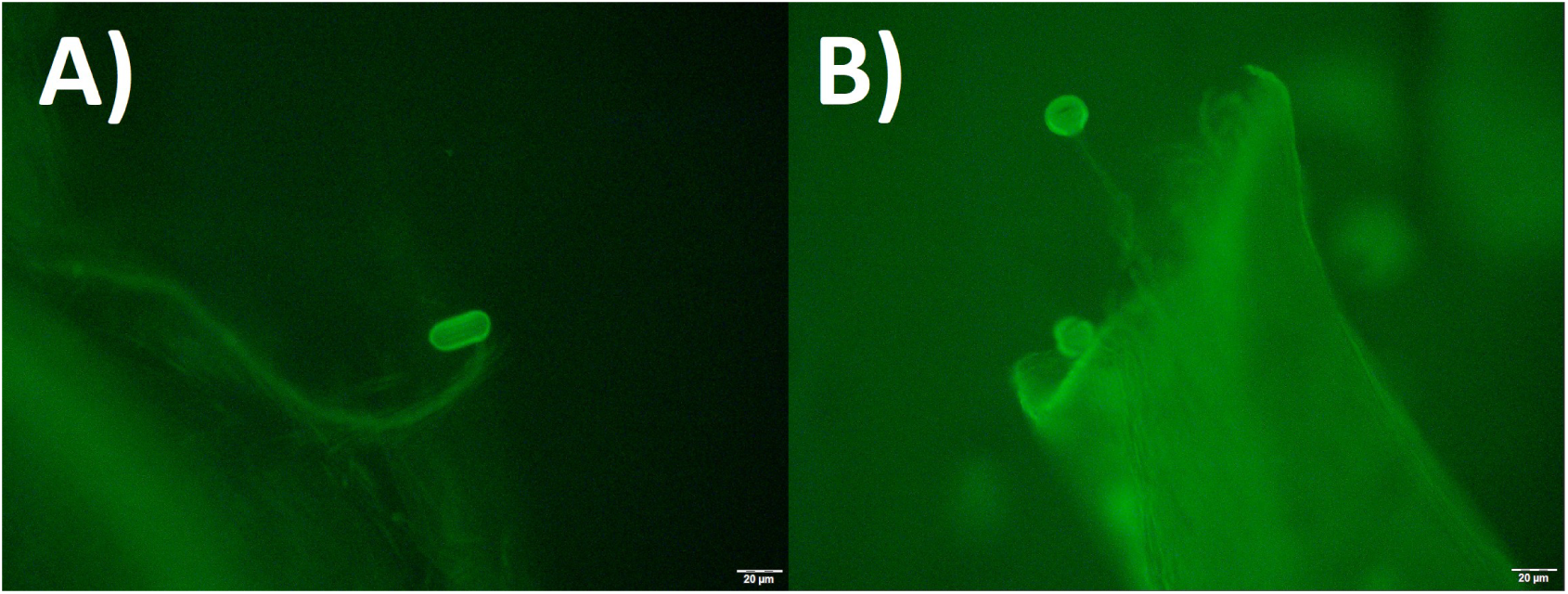
Pollen grains and tubes in the pistils: Images from a fluorescence microscope of *A. ilicifolius* (A.) and *A. officinalis* (B.).

### 2.5. Fruit and seed collection

Fruits were only collected from *Acanthus ilicifolius* from each site. *A. ilicifolius* starts flowering in March and it’s fruits were available at the time of our fieldwork. Fruit production of *A. ilicifolius* was assessed as the number of fruits collected per infructescence (3-12 infructescences per site) and seed production was estimated by counting the number of seeds per fruit in each infructescence. The number of seeds per fruit ranges from zero up to four in a fully seeded pod. Fruiting of *Avicennia officinalis* in that area occurs during July-August, which is at the peak of the rainy season when the forest is inaccessible and fruit collection was thus not feasible for every site. Local collectors were unable to reach sites inside deep forest due to the high water level and unavailability of transport.

### 2.6 Statistical analysis

Shannon’s Diversity Index (Shannon, 1948) was used to compare plant and pollinator species diversity between the sites. We analysed the impact of the position of the sites along the gradient from highly disturbed to the least disturbed parts of the forest (expressed as the distance of the sites from the village in km) on plant abundance and diversity using generalized linear models (GLM). We used Gaussian error distribution for plant species richness and Shannon’s diversity index, and binomial error distribution with overdispersion (“quasibinomial”) for proportion of plant cover. Multiple values of flower abundance, insect visitation, and pollen grains deposited on stigmas were measured repeatedly at each site, so we used generalized linear mixed-effects model (GLMM) with site identity as a random factor and Poisson error distribution for these response variables. We also used the duration of the observation period and the number of flowers in the observation window (both log-transformed) as an offset in the GLMM of insect visitation to properly analyse variation in visitation per flower per hour (Reitan & Nielsen, 2016). We also performed a redundancy analysis (RDA) to test for changes in the composition of the flower visitor assemblages with the increasing distance from the village area, separately for the two plant species. Finally, we analysed fruit and seed production in *A. ilicifolius* using similarly constructed GLMMs, with the number of flowers in an inflorescence used as an offset (log-transformed) when analysing the number of fruits per flower, and the number of fruits used as an offset (log-transformed) when analysing the number of seeds per fruit. We used R 3.4.4. (R Core Team 2018) for all analyses and plots; GLMMs were fitted using the lme4 package (Bates *et al*. 2015).

## 3. Results

All data underlying the results and supplementary materials are available in a Figshare repository: https://doi.org/10.6084/m9.figshare.11877615.

### 3.1 Plant diversity and abundance

Overall, we found 13 plant species of 9 families in the sampled sites (data: https://doi.org/10.6084/m9.figshare.11877615). However, more plant species can be found in the wider area.

We observed the lowest plant species richness and diversity in the forest patches nearby the village (Fig. 4A and 4B), which were dominated by mostly *Sonneratia apetala* Buch-Ham., *Excoecaria agallocha* L., and *Avicennia officinalis* L., all of them are tree species. Plant species richness and diversity increased along the gradient from the village towards the undisturbed forest interior (GLM, F=6.4, P=0.030 for species richness, and F=4.5, P=0.060 for Shannon’s diversity index). *A. ilicifolius* was the only shrub plant in the forest patches in the village area. The understory plant cover increased significantly towards the forest interior (GLM, F=29.6, P=0.0003; Fig. 4D), unlike the canopy cover (GLM, F=3.4, P=0.096; Fig. 4C)

**Figure 4:**
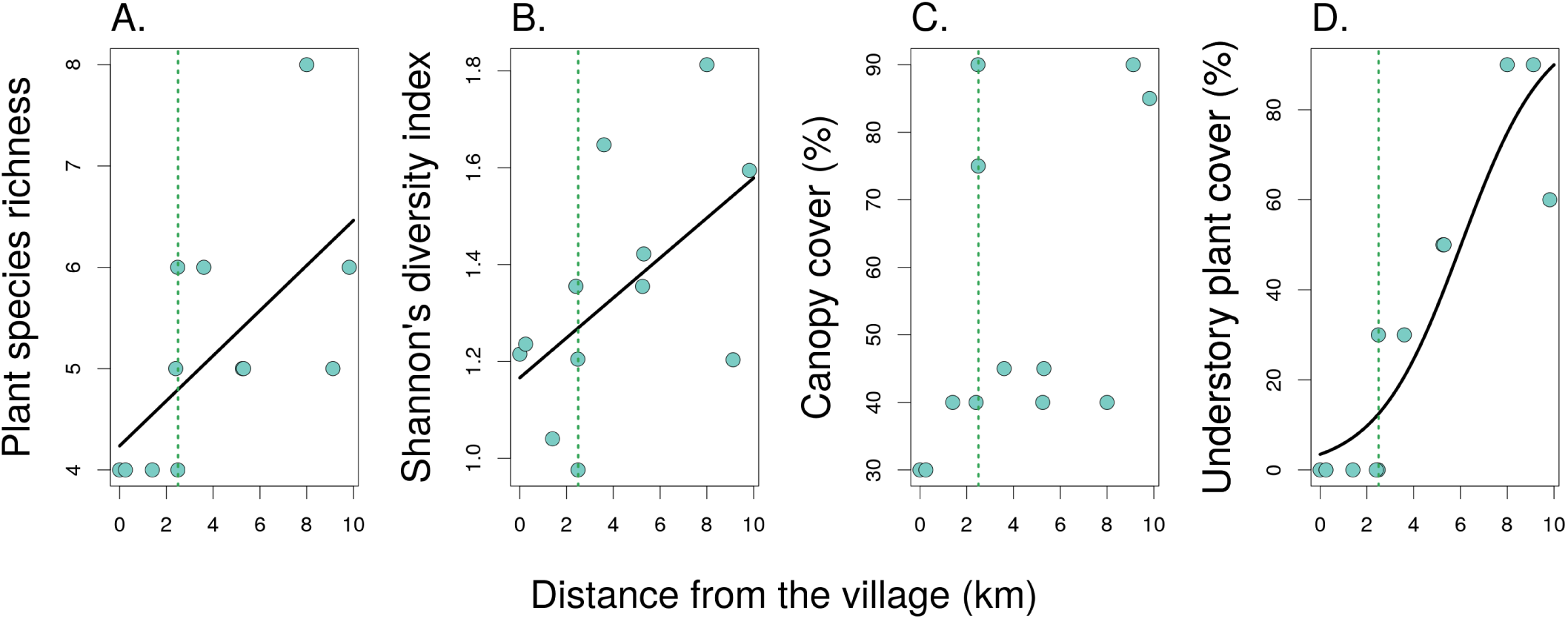
Plant diversity and abundance along the disturbance gradient and distance from the village towards the pristine forest: plant species richness (A.), Shannon’s diversity index (B.), canopy cover (C.) and understory plant cover (D.) for each site.

We also estimated the plant cover individually for our two target plant species and counted the number of flowers/m^2^ for both plant species to assess their floral abundance. Plant cover of the shrub *A. ilicifolius* gradually increased from the village towards the forest interior (GLM, F=56.3, P<0.0001; Fig. 5A) but did not change significantly in case of *A. officinalis* (GLM, F=0.32, P=0.58; Fig. 5B). Flower density of *A. ilicifolius* did not vary significantly along the gradient (GLMM, X^2^=0.95, P=0.33; Fig. 5C), while *A. officinalis* showed decreasing flower density from the village towards the deep forest (GLMM, X^2^=8.9, P=0.0029; Fig. 5D).

**Figure 5:**
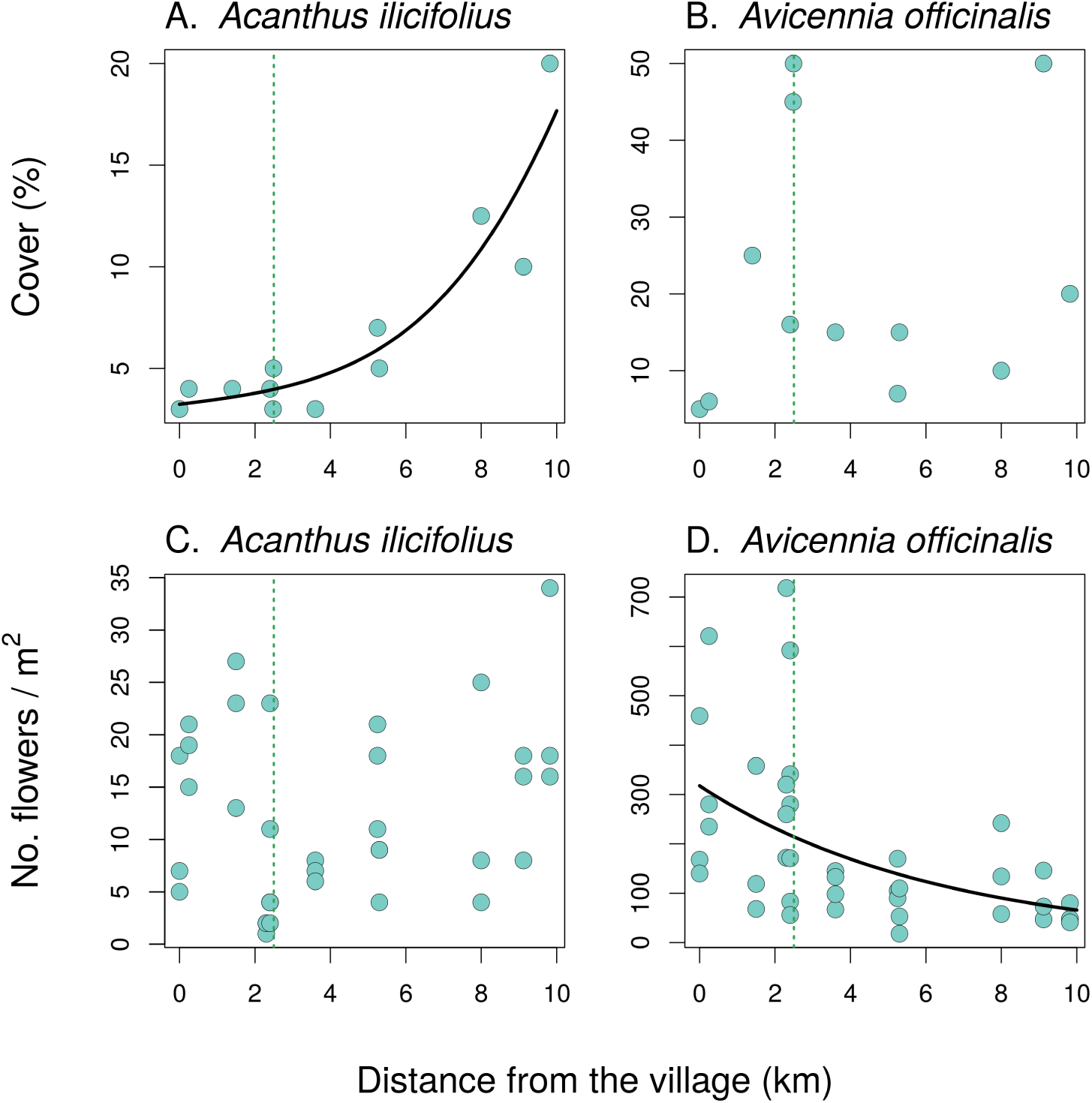
Individual plant cover and number of flowers for two target species: **A**. Plant cover for *A. ilicifolius* (shrub). **B**. Plant cover for *A. officinalis* (tree). **C**. Flower abundance for *A. ilicifolius* and **D**. Flower abundance for *A. officinalis*. X-axis showing the distance of the sites along the gradient from the village towards the undisturbed forest interior (**A**-**D**).

### 3.2 Insect diversity and flower visitation rate

Flower visitor community in our sampling period of this part of the forest consisted of the major insect groups, such as Hymenoptera, Diptera, Coleoptera and Lepidoptera. We observed total 4431 pollinating insects and randomly collected total 536 pollinating insects excluding three *Apis* species some easily recognised insects. We identified 105 insect species or morphospecies from at least 27 families (list of species: https://doi.org/10.6084/m9.figshare.11877615). Hymenoptera made up to 80% of the total number of pollinator individuals. Among them, bees were the biggest group and around 44% of them belonged to the genus *Apis. A. dorsata* Fabricius, 1793, also known as the giant honey bee, was the most abundant overall both in the village and inside the forest. Additionally, *Apis cerana* Fabricius, 1793, the eastern honey bee or the Asiatic honey bee, and *Apis florea* Fabricius, 1787, the dwarf honey bee, were found in the forest patches near the village but were completely absent in the deep forest. Next to the three honey bee species, solitary bees were the major insect groups among pollinators, consisting 38 species from 7 families. Wasps from the family Vespidae were the most diverse insect family with 22 morphospecies. Among the non-bee pollinators, flies, beetles and butterflies made up to 15% of the total pollinators. No bird was observed as pollinator for *A. ilicifolius*.

We found no significant changes in the total visitation of the two plant species by pollinators along the gradient from the village to the undisturbed forest interior (GLMM, X^2^=1.5, P=0.22 for *A. ilicifolius*, and X^2^=2.8, P=0.096 for *A. officinalis*). In addition, there were no significant changes in species richness of pollinators along the gradient for both plant species (GLM, F=0.49, P=0.050 in *A. ilicifolius* and F=1.4, P=0.26 in *A. officinalis*). The Shannon’s diversity index of the pollinator community also did not change in *A. ilicifolius* (GLM, F=2.1, P=0.17) and *A. officinalis* (GLM, F=1.5, P=0.25). However, species composition of the pollinators varied along the gradient as revealed by the redundancy analysis (RDA). The distance of the sites from the village explained 21.36% of total variance in species composition in the flower visitors of *A. ilicifolius* (RDA, F=2.72, P=0.0078) and 23.83% in A. *officinalis* (RDA, F=3.13, P=0.0021) with some species or groups more abundant in the sites close to the village area (e.g. *Apis cerana, A. florea*, and Diptera) and others (*A. dorsata* and Coleoptera) in the forest interior (Fig. 6, Fig. 7, Fig. 8, Table 1).

**Figure 6:**
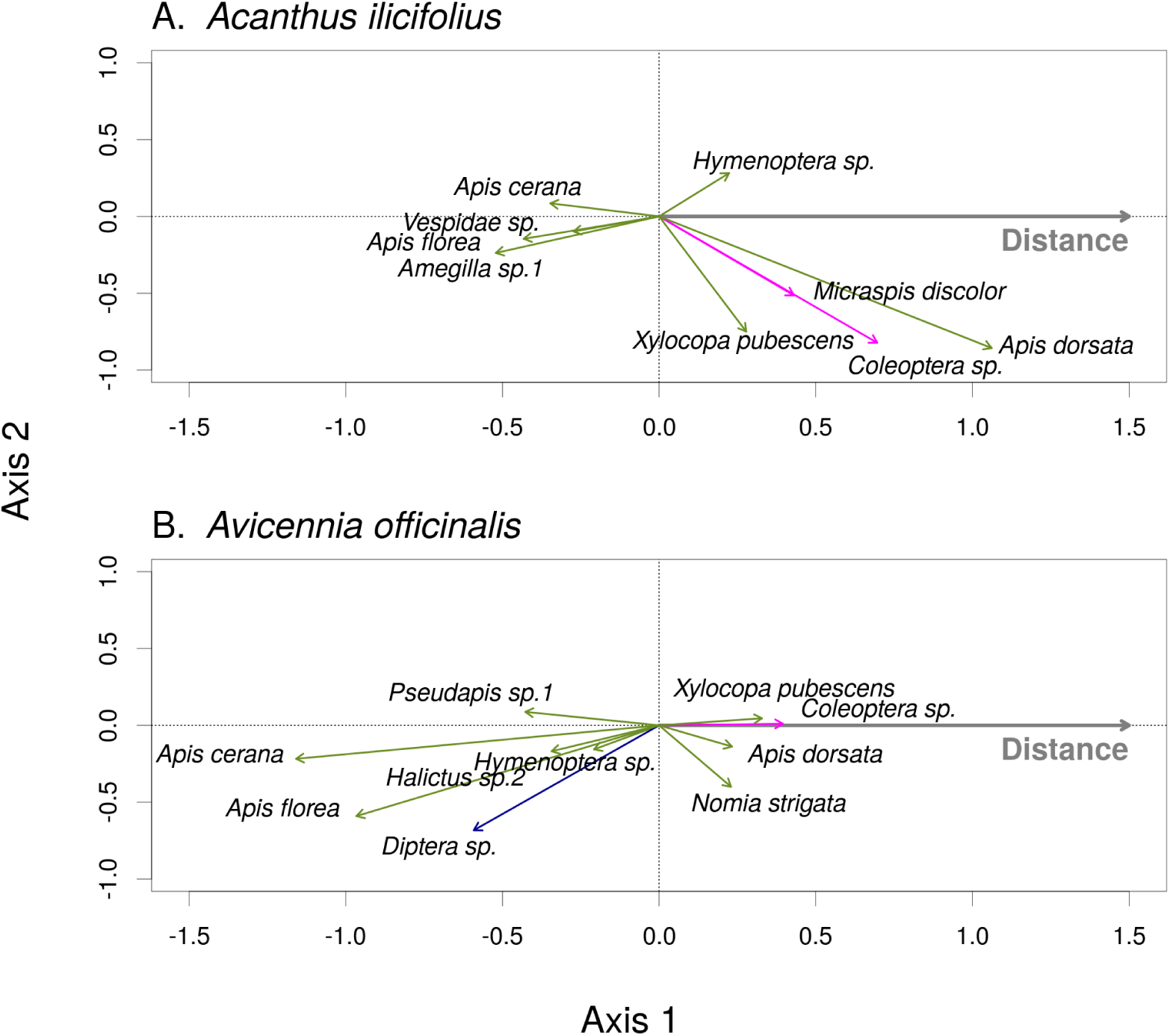
The composition of flowers visitors of *Acanthus ilicifolius* (A.) and *Avicennia officinalis* (B.) changed along the forest disturbance gradient. Results of RDA which show how abundance of individual flower visitor species on the two plant species changed with increasing distance from the village. Hymenoptera are displayed by green arrows, Coleoptera by magenta, and Diptera by blue arrows. Species whose abundance was little affected by the distance from the village (species with scores on Axis 1 <|0.2|) are not shown for clarity. Species with arrows pointing to the left were associated mostly with the fragmented forest close to the village, while species with arrows pointing to the right were found mostly in the deep forest far from the village.

**Figure 7:**
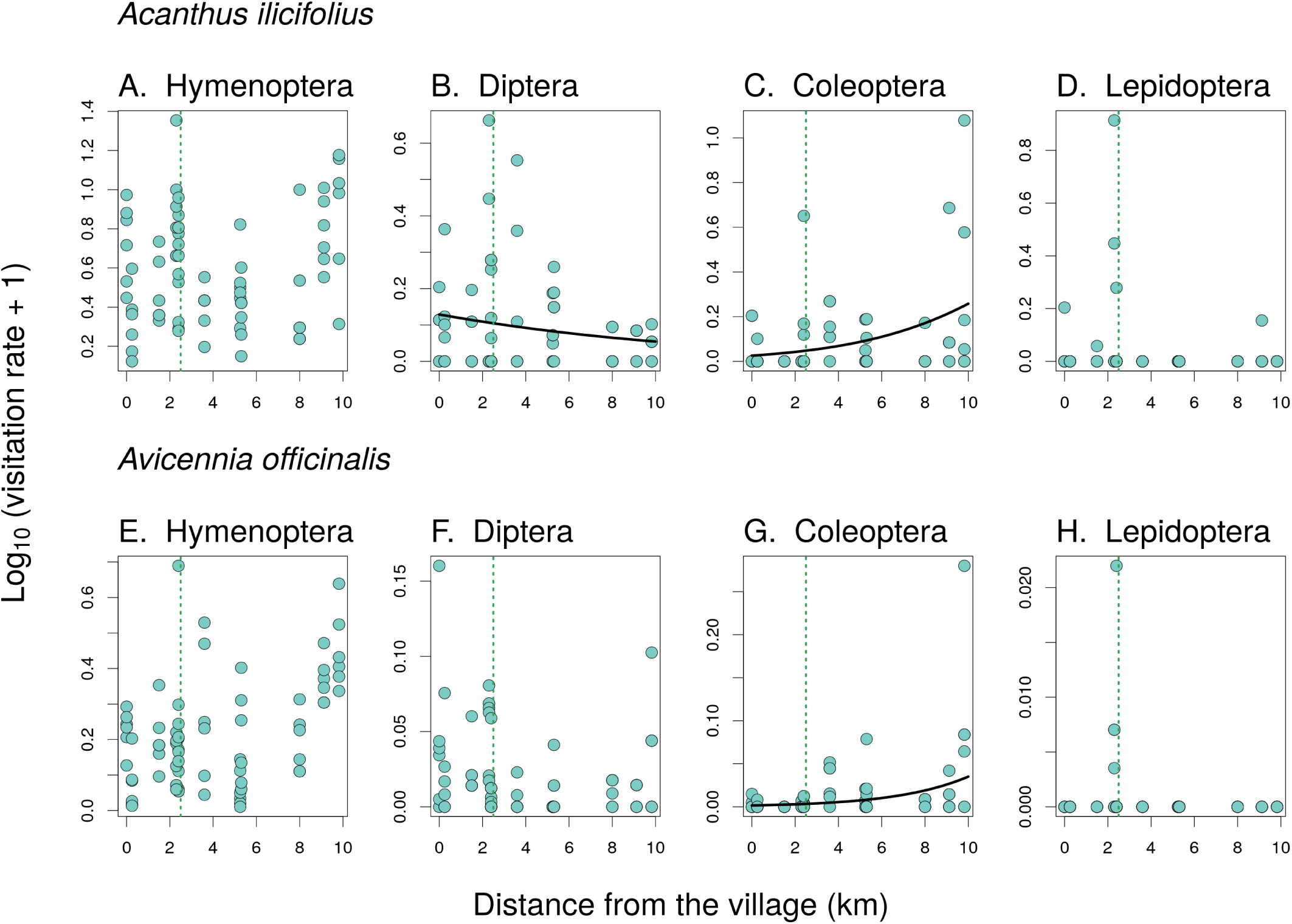
Visitation rate of insect orders on the two plants. The number of visits per flower and hour on *Acanthus ilicifolius* (A. - D.) and *Avicennia officinalis* (E. - H.). The estimated relationship is plotted as a line only in cases where it was statistically significant according to a likelihood ratio test (see Table 1). The vertical dotted green line shows the point along the disturbance gradient where the continuous forest begins and continues further away from the village.

**Figure 8:**
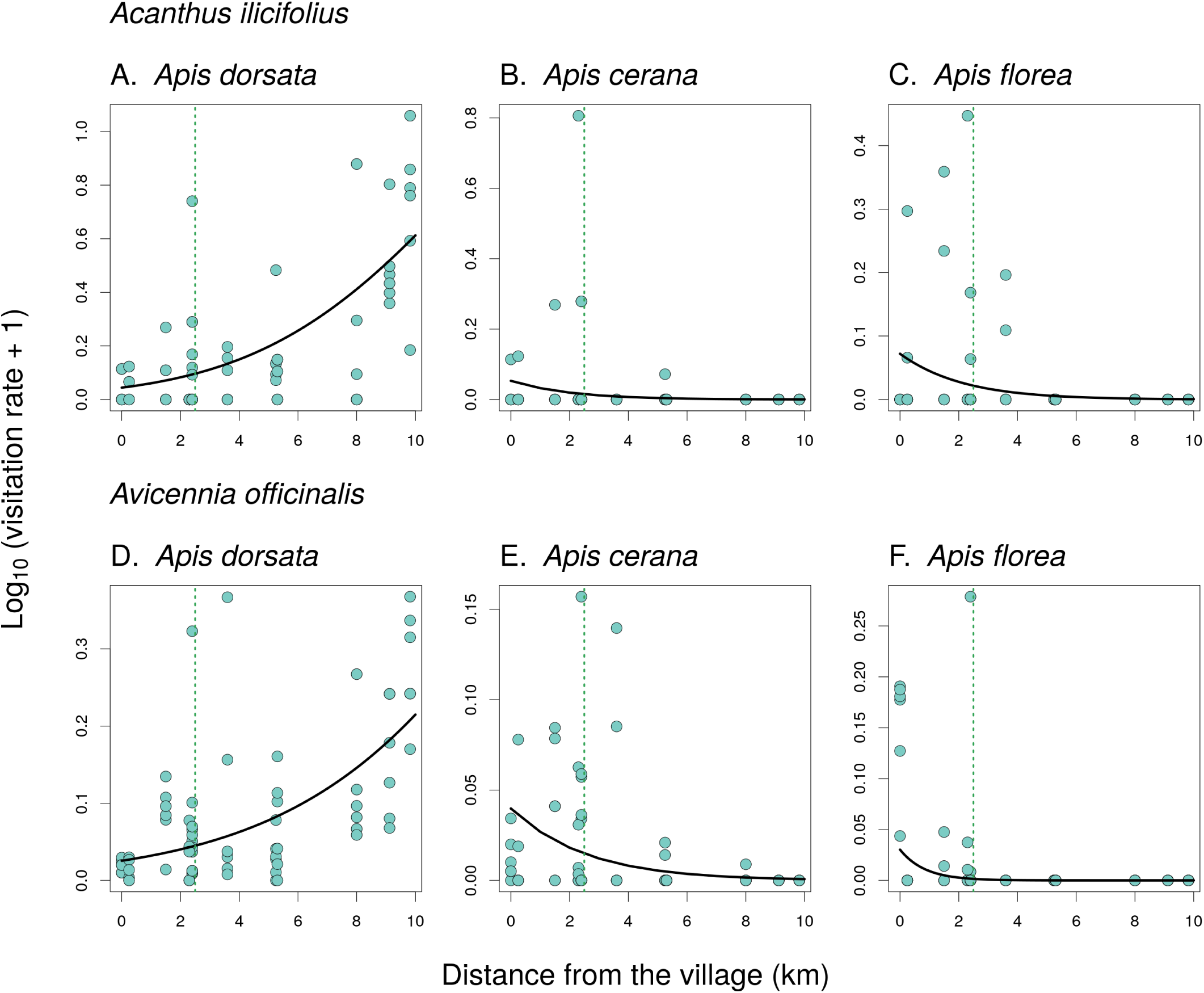
Visitation rate of the three species of honey bees on the two plants. The number of visits per flower and hour on *Acanthus ilicifolius* (A. - C.) and *Avicennia officinalis* (D. - F.). The relationship between the visitation rate and distance from the village was statistically significant in all cases according to a likelihood ratio test (see Table 1). The vertical dotted green line shows the point along the disturbance gradient where the continuous forest begins and continues further away from the village.

Both plants were visited mostly by Hymenoptera at a rate which did not vary along the disturbance gradient, while the visitation rate by Diptera on *A. ilicifolius* decreased and visitation rate by Coleoptera increased along the gradient on both plants (Table 1). Lepidoptera was observed rarely and mostly at sites along the edge of the continuous portion of the forest (Fig. 7). The three honey bee species of the genus *Apis* were the most frequent visitors on flowers of both plants, but they responded differently to the disturbance gradient (Fig. 8). *A. cerana* and *A. florea* abundances decreased with the distances of forest patches from the village towards the forest, while the number of *A. dorsata* increased gradually with the increasing distances of forest patches from the village (Fig. 8).

### 3.3 Pollen deposition and seed production

Although we observed significant variation in the composition of pollinator communities of both plants along the disturbance gradient from the village to the forest interior, pollination was not highly affected by these variations. The number of pollen tubes in the *A. ilicifolius* did show a significant increase in the total number of pollen grains deposited on its stigmas with the distance from the village towards the forest interior (GLMM, X^2^=4.2, P=0.041; Fig. 9A.), but this did not translate into differences in fruit and seed production. That is, the number of fruits per infructescence was not affected by the distance along the gradient from the village towards the forest interior (GLMM, X^2^=0.29, P=0.59), the same holds for the number of seeds per fruit (GLMM, X^2^=0.069, P=0.79). The second species, *A. officinalis*, showed no differences in the total number of pollen grains deposited per flower along the disturbance gradient (GLMM, X^2^=1.3, P=0.25; Fig. 9B.), while seed set data were not available.

**Figure 9:**
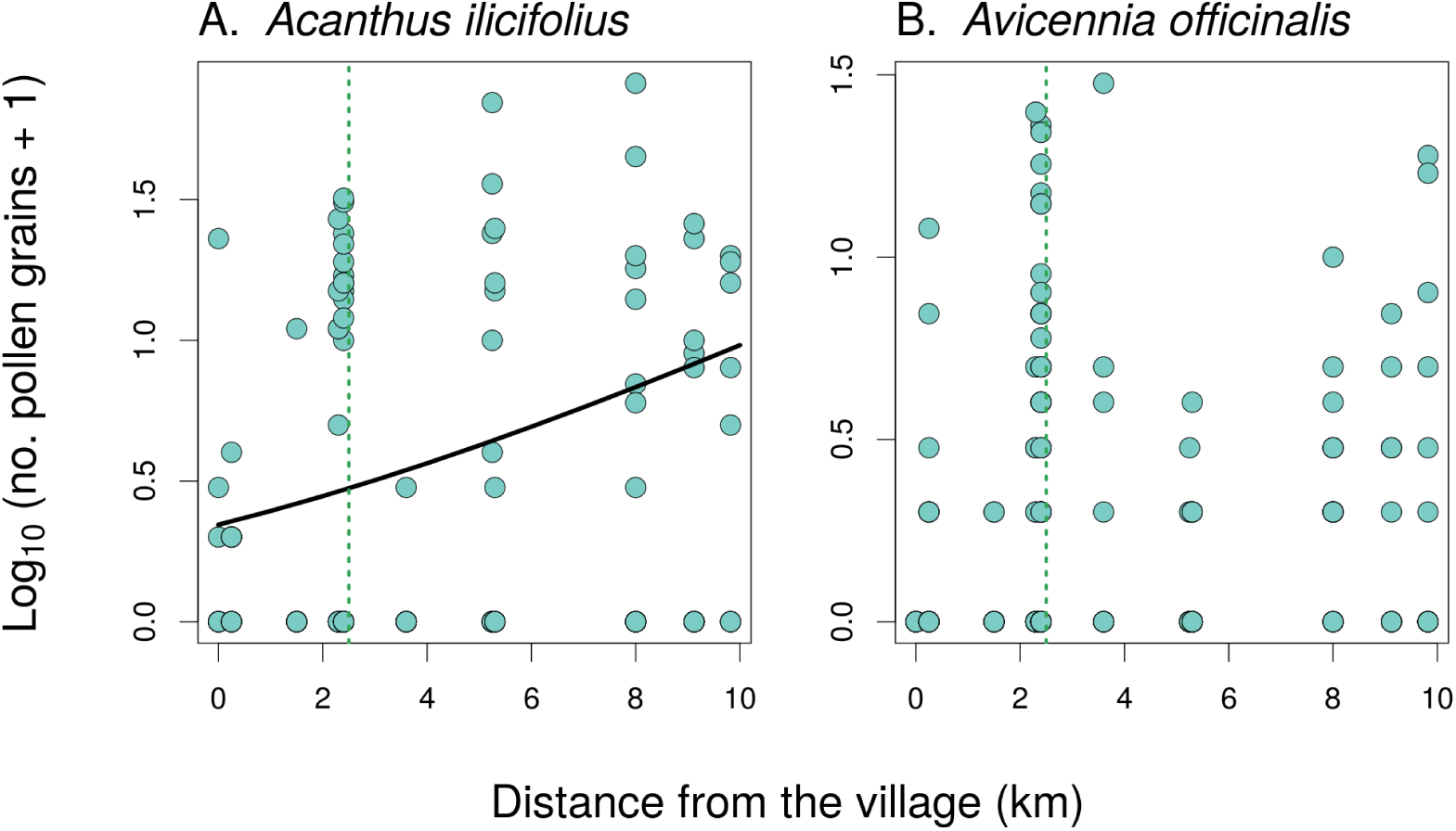
Pollen deposition on stigmas of the two plant species. The number of pollen grains deposited on stigmas of *A. ilicifolius* (A.) and *A. officinalis* (B.) in relation to the distance from the village towards the forest interior. The relationship was statistically significant only in *A. ilicifolius*. The vertical dotted green line shows the point along the disturbance gradient where the continuous forest begins and continues further away from the village.

## 4. Discussion

### 4.1 Impact of human disturbances on plant diversity, pollinator communities and pollination

Our research shows changes in the plant community structure, composition of the pollinator communities, and flower visitation patterns along the gradient of decreasing human disturbance from village area towards a relatively pristine forest interior. Forest patches nearby the village had the lowest number of plant species and the lowest values of the Shannon’s diversity index. These fragmented forest patches are used by the local people as grazing land for their domestic animals and plant leaves and stems were regularly collected for fuel and fodder and intentionally kept clear to deter tigers as they prefer to hide in the bushes for hunting (Badhwar, 1988). Furthermore, pollutants from boats and households may hamper the regrowth of plants in such forest patches (Santos *et al*., 2012) and resulted to have almost no understory and very low plant species richness and diversity. Likewise, *A. ilicifolius* was lower in the patches nearby the villages and increased significantly towards the deep forest. Although plant cover increased with the distance from the village, flower production per unit area did not show any differences. Contrastingly, the percentage cover of *A. officinalis* did not show any change along the disturbance gradient but its flower abundances decreased towards the deep forest. This can be due to the rain deficiency in this part of the forest as *Avicennia* species flowering time shows a high sensitivity to rainfall (Opler *et al*., 1976; Reddy *et al*., 1995) or due to the increasing salinity as several studies showed delayed flowering in plants due to the increasing salinity (Maas and Poss, 1989, Khatun and Flowers, 1995). This forest is lying in the Bay of Bengal delta and with the advantage towards the deep forest from the village leads to higher salinity as the sea gets closer (Haque and Reza, 2017). We also noticed that *A. officinalis* had an unusually high cover in some of the patches at the transition between the village and continuous forest due to the partial plantation by the forest department to support the restoration of the forest (Saenger and Siddique, 1993; Rahman and Rahman, 2015).

The giant honeybee (*A. dorsata)* was the major pollinator for both species in every site with a sharp increase from the village towards the forest interior. Although *A. dorsata* was reported to be a vital pollinator for both cultivated crops and wild plants (Robinson, 2012), they usually forage in more abundant flower resources (Punchihewa *et al*., 1985) and may be a poorer competitor than the other two *Apis* species (Koeniger and Vorwohl, 1979). Studies showed that our three honey bee species compete for food, with *A. florea* and *A. cerana* being the stronger competitors than *A. dorsata*, and this competition can be avoided by differentiation of foraging (Koeniger & Vorwohl, 1979). This may explain why the abundance of *A. dorsata* is lower in forest patches nearby the village and higher in the deep forest as *A. cerana* beehives were located in the village patches and *A. florea* was only present in the village sites. It has been reported that the presence of domesticated *A. cerana may* affect the abundance of *A. dorsata* in human disturbed areas (Samejima *et al*., 2004). Moreover, forest patches nearby the village are more exposed to both professional and non-professional honey collectors and naturally occurring *A. dorsata* hives are frequently disturbed, extracted, and even destroyed by the honey-collectors which may lead to low number of hives in the village areas. On the other hand, *A. cerana* is completely domesticated in that area and they were able to forage both in the forest and village patches within their foraging distance (Partap, 2011) while they were absent in the deeper forest. Among the three species, *A. florea* has a distinct habitat preference and was only found in village areas. This smaller bee prefers to build their nest in lower branches, in sunny location (Whitcombe, 1984) and forest patches near the village offer more suitable nesting sites in terms of their habitat preference, compared to the deeper mangrove forest.

*A. florea* tend to swarm and transfer nest swiftly and prefer to stay close to the abundant food and habitat resources (Whitcombe, 1985). Unlike other *Apis* species, this species does not migrate when the flower resources are scarce and shortage of their flight range make them more aggressive towards other bees but generally niche compartmentalization between the flower resources would minimize the competition (Koeniger & Vorwohl, 1979) and different studies on the *Apis* species showed that these three species can co-occur without any significant competition (Punchihewa, *et al*., 1985, Oldroyd, *et al*., 1992, Koetz, 2013). However, interactions between the domesticated *A. cerana* and other pollinators are not well-known. A number of recent studies on European honeybees *Apis mellifera* showed their strong negative effects on wild pollinators (Magrach *et al*. 2017, Henry & Rodet 2018, Hung *et al*. 2019). Exploring the interactions between the Asian honeybees *A. cerana* and wild pollinators in similar detail will thus be an important topic for future research.

Although three species of *Apis* made up half of the pollinating insects for both targeted plants, solitary bees played the second most important role in visiting the flowers. The overall abundance of Hymenoptera increased towards the deeper forest for *A. ilicifolius* but decreased for *A. officinalis*, but this can be the result of decreased flower abundance of this species. On the other hand, the abundance of *Xylocopa pubescens* increased towards the deep forest for both plant species, although their abundance was not as significant as other pollinators, despite their well established role as a pollinator for *A. ilicifolius*. We did not observe any birds visiting *A. ilicifolius*, although Primack and Tomlinson (1980) reported sunbirds as pollinator for this species in Australia. Unlike *Apis* spp., our study found that the disturbance gradient had little effect on the total flower visitor abundances and diversity of this group. Although some studies suggested that the species richness and population density of solitary bees may decrease proportionally with the increasing human disturbances (Inoue *et al*., 1990; Liow *et al*., 2001), another study showed the opposite where wild be communities were reported to be persistent against the human disturbances or even to benefit under particular circumstances (Stein *et al*., 2018). However, we lack detailed information about the biology and foraging behaviour of individual species, apart from *Apis* spp. discussed above, which prevents more detailed assessment.

### 4.2 Perspectives for forest conservation

Many plants in the mangrove forest are dependent on the insect pollination and similarly, mangrove provides an excellent forage for bees (Lacerda, 2002) and other insects. Human disturbance impacts on both plant and animal diversity are likely to be severe, therefore, we need to focus on developing sound conservation policies for the mangrove forests, such as the Sundarbans. Based on our results, it seems that changes in the composition of the pollinator community along the gradient of human disturbances did not affect the pollination success of the studied greatly, but the plant diversity and cover of the understory plants were significantly lower in patches close to the village. This suggests that despite the successful pollination and seed production, human exploitation interrupts forest regeneration and likewise affects the pollinator community. Moreover, overexploitation of the wild-living giant honey bees, *A. dorsata*, was likely responsible for the lower abundance of this species in forest patches near the villages compared to the pristine part of the forest. However, almost no information is available on how honey hunting affects the colony survival, growth and migration of *A. dorsata*. Continuous destruction of its nests and habitat may lead to further decline of the giant honey bee. Local extinctions of *A. dorsata* have been reported across their vast distribution range (Oldroyd and Nanork, 2009) and deserve attention regarding their conservation. On the other hand, based on the population growth rate and the rate of harvesting of *A. florea*, the other wild-living honey bee species, it is unlikely that this species will be affected by human disturbances at the same rate as other honey bees (Oldroyd & Wongsiri, 2006). The third local honey bee species, *A. cerana*, is domesticated and kept by the local beekeepers and it is unlikely to go extinct. Studies in other low-land forest areas in Asia showed that *A. dorsata* immigrates into the forests in the mass flowering season and when the amount of floral resources drop, they leave the forest (Itioka *et al*., 2001). In contrast, the resident *Apis* species and other solitary bees, stay year around and pollinate the flowering plants for entire period (Sakai, 2002). Hence, keeping domestic honey bees in the mangrove areas is widely accepted as non-harmful from the conservation point of view. However, more intensive research will be needed to decide whether keeping domestic beehives in this area is beneficial or harmful for the local pollinator communities and plant diversity.

Although our results show that changes of the composition of the flower visitor community along the gradient of human disturbance in our case likely do not affect the reproduction of the studied plants, human activities negatively affect the mangrove forest in other ways, mostly by disrupting forest regeneration by clearing the understory. Also, we conclude that bees are the most important pollinators in this forest, but *Apis dorsata* is threatened by human activities, in particular by harvesting of its honey. The forest provides vital resources for the local people, so to prevent further deterioration of the state of the forest, it is necessary to initiate more intensive conservation approaches, e.g. mangrove tree plantation with a focus on rare species, and increase awareness about the necessity of mangrove conservation among the locals and involve them directly through the community based approaches (López-Portillo *et al*., 2017). While honey harvest is an important source of income for the local people, honey hunters should be encouraged and trained to harvest honey in a non-destructive sustainable way with proper equipment to minimise the impact on the bee hives (Purwanto *et al*., 2000; Waring and Jump, 2004).

## 5. Acknowledgements

We would like to thank students from the University of Chittagong and Khulna University, Bangladesh for their help in the field work and the technical support from both Universities. Our thanks to Jiří Hadrava and Martin Šlachta for help with species identification of Diptera and Hymenoptera. We also thank the Forest Department of Bangladesh for granting the permission and security to work in this forest. We are also grateful to Jana Jersáková for the help with laboratory analysis. This project was funded by National Geographic Society (Grant-EC-376R-18) and also partly supported by the Czech Science Foundation (GJ17-24795Y) and by the University of South Bohemia in České Budějovice, Czech Republic. The funders had no role in the study design, data collection, analysis and preparing the manuscript.

